# Cardiomyocyte mechanical memory is regulated through the talin interactome and DLC1 dependent regulation of RhoA

**DOI:** 10.1101/2023.07.19.549635

**Authors:** Emilie Marhuenda, Ioannis Xanthis, Pragati Pandey, Amar Azad, Megan Richter, Davor Pavolvic, Katja Gehmlich, Giuseppe Faggian, Elisabeth Ehler, James Levitt, Simon Ameer-Beg, Thomas Iskratsch

## Abstract

Mechanical properties are cues for many biological processes in health or disease. Likewise, in the heart it is becoming clearer that mechanical signals are critically involved in the disease progression. Cardiomyocytes sense the mechanical properties of their environment at costameres through integrins and associated proteins, including the mechanosensitive protein talin as an integral component. Our previous work indicated different modes of talin tension, depending on the extracellular matrix stiffness. Here, we wanted to study how this leads to downstream mechanotransduction changes, further influencing the cardiomyocyte phenotype. Combining immunoprecipitations and Fluorescence Recovery after Photobleaching (FRAP) experiments, we identify that the talin interacting proteins DLC1, RIAM and paxillin each preferentially bind to talin at specific extracellular matrix stiffness and this interaction is preserved even in absence of tension. This demonstrates a mechanical memory, which we confirm further *in vivo* in mouse hearts. The mechanical memory is regulated through adhesion related kinase pathways. Optogenetic experiments using the LOVTRAP systems confirm direct competition between the individual proteins, which again is altered through phosphorylation. DLC1 regulates RhoA activity in a stiffness dependent way and both loss and overexpression of DLC1 results in myofibrillar disarray. Together the study demonstrates a mechanism of imprinting mechanical information into the talin-interactome to finetune RhoA activity, with impacts on cardiac health and disease.

## Introduction

Cardiomyocytes are the contractile cells in the heart and their proper function is regulated through a complex signalling network that includes electrical, chemical and also mechanical signals^1^. The mechanical signals again are varied and include e.g. pressure and stretch from filling of the heart with blood, but also importantly the sensing of the stiffness of the extra cellular matrix. The latter is changing during development. Importantly it is also changing during ageing and in heart disease, where excessive crosslinking of collagen through lysyl oxidases (LOX) and LOX like enzymes can lead to a stiffening of the heart, cardiomyocyte phenotypic changes and heart failure with preserved ejection fraction (HFpEF) ^2-9^. Cardiomyocytes attach to the extracellular matrix (ECM) through so-called costameres, rib-like structures at the level of the sarcomeric Z-disc, that contain integrins as well as other multi-molecular complexes for adhesion (e.g. the dystrophin glycoprotein complex) and/or signalling^10^. The cardiomyocyte integrin adhesion has many of the proteins that are also found in focal adhesions, including talin and vinculin which attach to cytoplasmic actin. The cytoplasmic actin in return, is further connected to the sarcomeric Z-disc through actin crosslinkers such as α-actinin and plectin ^11^.

Previous studies, including our own work documented the role of ECM stiffness in determining the contraction dynamic and contractile work of cardiomyocytes^12-17^. Moreover, our own previous work suggested that cardiomyocytes can sense the ECM stiffness at integrin adhesions, through a combination of myofibrillar and non-myofibrillar contractions^17^. Non-muscle myosin, which contracts the cytoplasmic actin, generates a basic tension on integrin adhesions, which persists during diastole and to which the forces from the myofibrillar contractions are cyclically added.

Moreover, non-muscle myosin is increasingly localized at the costameres in heart disease. Depending on the stiffness of the ECM and the level of non-myofibrillar contraction this leads to different dynamics of tension on the integrin adhesion protein talin, which is unstretched on embryonic stiffness, experience cyclic tension on healthy adult heart stiffness or constant tension on fibrotic stiffnesses. The different downstream signalling influences cardiomyocyte function and potentially disease progression.

Talin is increasingly regarded as a key mechanical sensor^18-20^. It contains a head domain that binds to the integrins, as well as a rod domain, composed of 13 helical bundles (R1 to R13) that include actin binding sites and further bind to a substantial number of other proteins^18^. Importantly, all rod domains can unfold under physiological forces (between 5 and 12pN) and refold once tension is released^20^. This opening and closing not only can lead to a remarkable extension of the protein to over 800nm^20^, but also to opening of cryptic binding sites, most prominently for vinculin^21^, which can then lead to adhesion reinforcement. Indeed, our previous work suggested enriched localisation of vinculin to force transduction sites (in this case nanopillars that were being pulled by the cardiomyocytes) and this enrichment in localisation increased with pillar stiffness, suggesting adhesion reinforcement^17^.

However, apart from vinculin, other force dependent interactions have been reported for talin, including Rap1-Interacting Adaptor Molecule (RIAM, also known as amyloid beta precursor protein binding family B, or APBB1IP) which is binding only to the closed R3 domain and thus in a mutually exclusive way to vinculin^22^. Similarly, the molecule “Deleted in Liver Cancer 1” (DLC1) binds to the R8 domain only in the folded state, but dissociates once it is unfolded under force^23^. Indeed, DLC1 is an attractive candidate for downstream mechanosignalling in cardiomyocytes. DLC1 is located at the plasma membrane and adhesions, it acts there as a Rho GTPase activating protein (RhoGAP, i.e. negative regulator of RhoA activity)^23-25^. RhoA has been associated with cardiomyocyte hypertrophy and increased fibrotic response to chronic pressure overload^26-28^. On the other hand RhoA is essential for early heart development^29^ and has protective effects after cardiac injury, promoting cardiomyocytes survival^30^ and delaying the transition to heart failure^26^. Together this suggests the need for a tight spatial and temporal regulation of RhoA activity in response to diverse chemical (e.g. downstream of hypertrophic stimuli acting on GPCRs, like Phenylephrine, Endothelin-1 or Angiotensin II)^31^ or mechanical signals and DLC1 binding to talin would ideally couple the regulation to the mechanosensing machinery.

Here, we investigate the cardiomyocyte mechanosignalling and focus on stiffness dependent talin interactions. We find that DLC1 is a major RhoGAP at cardiomyocyte adhesions and consequentially, a knock down of DLC1 results in stiffness dependent cytoskeletal disruptions and alteration of cardiomyocyte mechanics. Strikingly, we find that stiffness regulates differential binding of the three talin R8 tail domain binding partners DLC1, RIAM, and Paxillin. The interaction is preserved even after tension is released (evidenced by co-immunoprecipitation experiments), indicating mechanical imprinting. We demonstrate the imprinting is regulated by integrin-related phosphorylation pathways (especially FAK and Src), as well as a hierarchical competition, which we confirm using optogenetic experiments using the LOVTRAP system^32^. In summary, our data suggests a mechanism of mechanotransduction and mechanical memory that depends on talin stretching and altered talin interactions. Overall, we find that mechanical imprinting modifies talin interactions through integrin related kinase signalling and competition for binding to the talin R8 domain. Together this determines cardiomyocyte function through regulating the levels of active RhoA, via DLC1 and could have strong implications on pathological cardiac remodelling.

## Results

### DLC1 is the major RhoGAP in cardiomyocyte adhesions

We previously found that cardiomyocytes sensing of extracellular matrix stiffness resulted in different modes of talin stretching (no stretching vs cyclic vs static)^17^. Because of its mechanosensitive binding to talin and its role in regulating RhoA^23^, we hypothesised that DLC1 could be involved in the downstream signalling. To test if DLC1 binding to talin can regulate the mechanical signalling in the heart, we first confirmed its expression in cardiomyocytes. Indeed, single cell and single nuclei RNAseq from fetal and adult human hearts, as well as from postnatal mouse hearts all suggested that DLC1 is consistently expressed in cardiomyocytes at the highest level, compared to all adhesion localised RhoA, Rac or CDC42 GAPs or GEFs(Figure 1A, Suppl Fig 1A,B)^33^. Similarly, our bulk RNAseq dataset of control mouse hearts also indicated DLC1 as major adhesion localised small GTPase regulator in the heart (Fig 1B). In agreement with this we find adhesion staining in neonatal rat cardiomyocytes (Fig 1C) and costameric pattern in wild type mouse hearts (Fig 1D). Intriguingly, DLC1 is further upregulated in the MLP knock out mouse, a model for dilated cardiomyopathy, where we find further enrichment at the intercalated disc as well as ectopic expression pattern.

**Figure 1:**
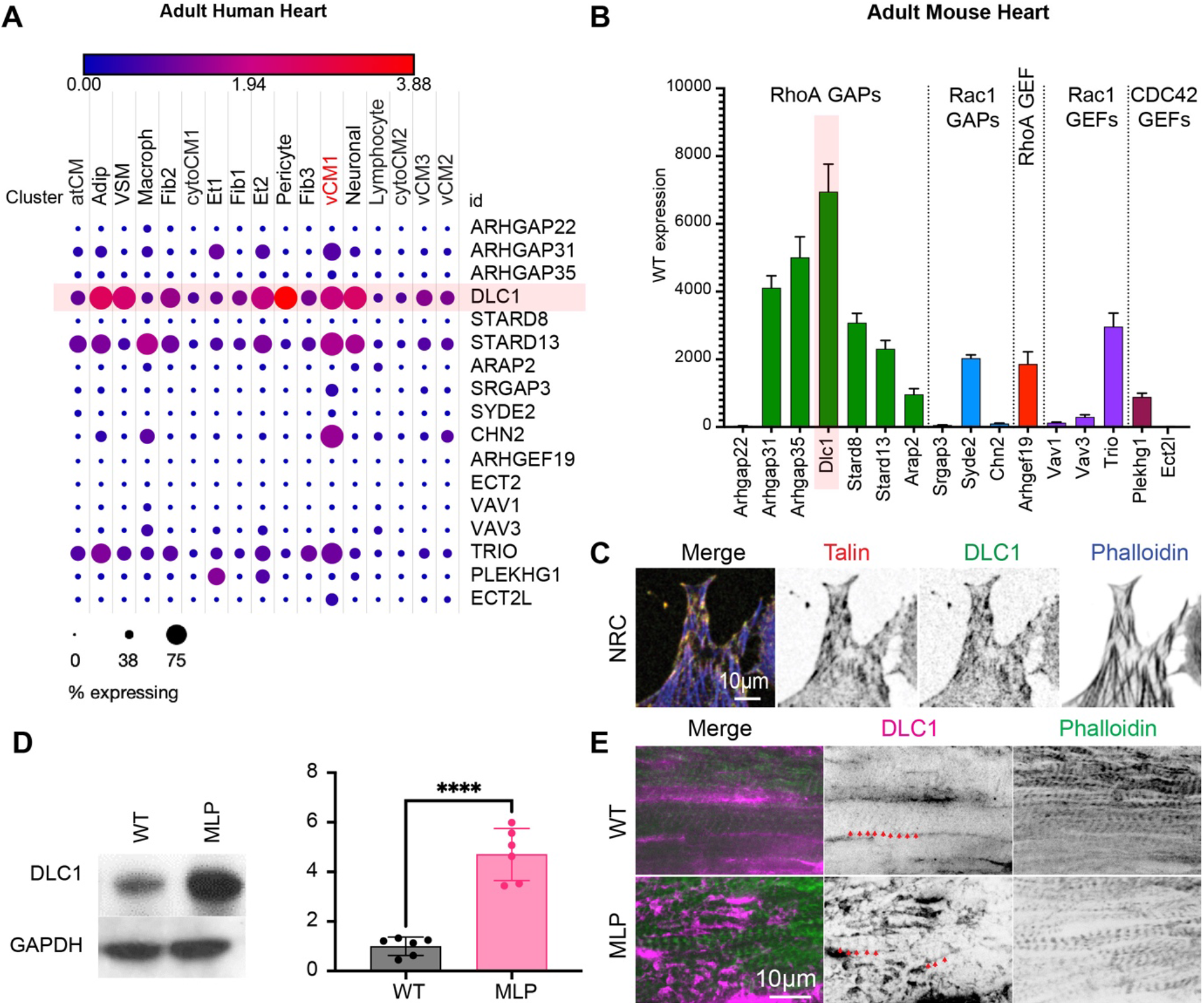
DLC1 is a major RhoGAP in cardiomyocytes. Expression of adhesion localised RhoA, Rac1 and CDC42 GAPs and GEFs (as identified by Müller et al ^33^) was analysed using the broad institute single cell portal in **A)** single nuclei RNA-Seq data of adult human hearts (287,269 nuclei from 4 women, 3 men, Age 39-60 years)^83^; atCM: atrial cardiomyocytes; Adip: adipocytes; VSM: vascular smooth muscle cells; Fib1,2,3: fibroblast populations; Et1,2: endothelial cell populations; vCM1,2,3: ventricular cardiomyocyte populations; cytoCM1,2: cytoplasmic cardiomyocyte populations. **B)** The single cell data is further consistent with our bulk sequencing analysis of adult wild type mouse hearts (four wild type mouse hearts)^84^; **C)** Immunostaining confirms the adhesion localisation in neonatal rat cardiomyocytes (NRC). **D)** DLC1 expression is increased in the MLP knockout mouse, as model for dilated cardiomyopathy. n=6 hearts for wt and the MLP knockout. **** p<0.0001 from unpaired t-test. **E)** Costameric staining is found in the healthy mouse heart (red arrow heads). Additionally, intercalated disc and ectopic expression is found in the MLP heart.

### DLC1 binds to Talin in a stiffness dependent way

DLC1 binds to the Talin R8 domain, which is atypical in the sense that it is nested within the R7 domain and requires higher forces for unfolding, albeit still within the physiological range^20^.

Additionally, RIAM and Paxillin bind to an overlapping binding site^34^. Paxillin is well described as a costameric protein in the heart and interacts amongst others with focal adhesion kinase (FAK)^35, 36^. Similar to DLC1, RIAM is expressed at costameres, whereby expression is increased MLP KO mice and human heart disease (Supplementary Figure S2). Because we previously found that talin is differentially stretched in cardiomyocytes plated on stiffness mimicking the embryonic, healthy adult and diseased heart^17^ we first tested how this different mechanical response would affect the interactions with DLC1, RIAM and Paxillin. For this we decided on a co-immunoprecipitation strategy with an anti-talin antibody (Figure 2A,B). Intriguingly, we found a strong stiffness dependence of the R8 domain interactions, whereby each protein had a specific optimal stiffness for interacting with talin: Paxillin interacted strongest at embryonic stiffness (1kPa), DLC1 interacted strongest at healthy adult heart stiffness (6kPa) and RIAM interacted strongest at elevated stiffness (20 and 130kPa).

**Figure 2:**
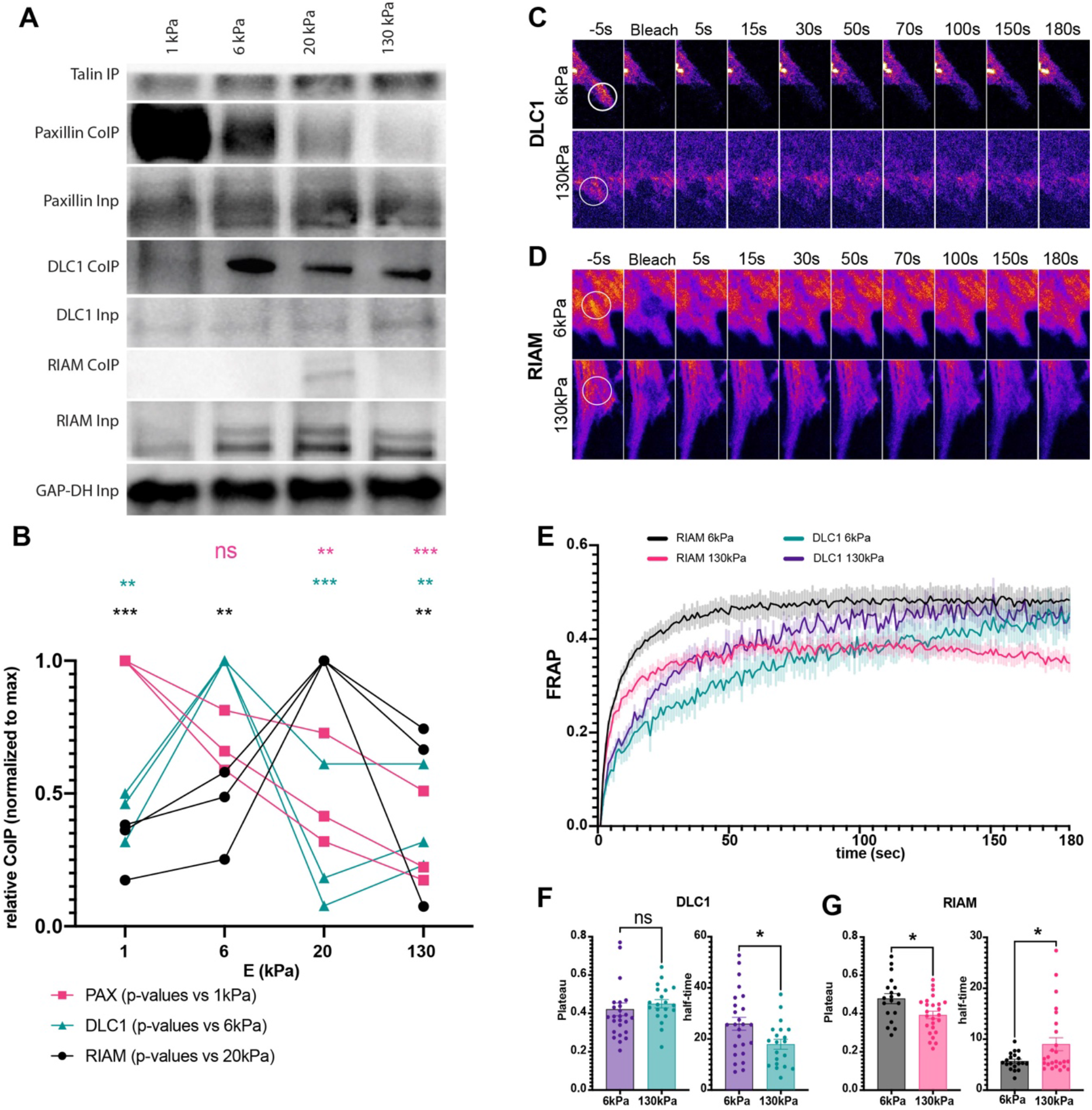
DLC1 and RIAM are binding to talin in a stiffness dependent way. **A)** Co-immunoprecipitation analysis for talin interactions. Neonatal rat cardiomyocytes were cultured on PDMS surfaces with the indicated stiffness for 7 days, before immunoprecipitation with an anti-Talin antibody. Samples were then subjected to western blotting and probed with antibodies against paxillin, RIAM and DLC1, indicating preferential binding at 1kPa (Paxillin), 6kPa (DLC1) and 20kPa (RIAM); quantified in (**B**) from three independent repeats; **C-G)** Fluorescence recovery after photobleaching (FRAP) assays of DLC1 (C) and RIAM (D) in neonatal rat cardiomyocytes. **E)** average recovery curves (mean +SEM); **F)** Quantification of plateau (mobile fraction) and recovery half-time for DLC1 and **G)** RIAM. *p<0.0332, **p<0.0021, ***p<0.0002, ****p<0.0001; p-values from 2-way ANOVA with Dunnett correction for multiple comparisons (B) or unpaired t-test (F,G);

Because talin is in a tension free state during the immunoprecipitation, this result suggested that the competitive interactions might be regulated through other mechanisms, such as posttranslational modifications. Importantly, this way the information about the mechanical environment could be imprinted into mechanical memory^37^. To test for the state of talin-interactions *in situ*, we performed fluorescence recovery after photobleaching (FRAP) experiments to investigate the stiffness dependent dynamics of DLC1 and RIAM at adhesion sites (Figure 2C-G). While we note that DLC1 was reported to bind to other adhesion proteins, such as tensin^38^, and RIAM can also bind to the talin R3 domain^22^, we nevertheless found a slower turnover of DLC1 on healthy adult stiffness (i.e. a stronger binding to the adhesion sites at 6kPa) and vice versa a slower turnover of RIAM on a fibrotic stiffness of 130kPa (Fig 2E-G), in agreement with the data from the immunoprecipitation. The plateau was unchanged for DLC1 suggesting no change in the immobile fraction (i.e. the stable binding molecular pool that does not undergo exchange during the timeframe of observation), while RIAM displayed a slightly reduced plateau and hence a larger pool of stable binding molecules at 130kPa (Fig 2F-G).

### DLC1 and RIAM binding is regulated through FAK signalling

Because our data suggested the cardiomyocytes interaction with different ECM matrix stiffnesses were imprinted into the talin interactome, we wanted to test if this might be regulated through integrin related phosphorylation pathways, including Src family kinases, FAK, Rho Kinase (ROCK) and PKC. For this we pre-treated the cardiomyocytes on healthy adult heart stiffness (6kPa) with the respective inhibitors (PP2, FAK Inhibitor, Y27632 and Bisindolylmaleimide I) before cell lysis and performed the co-immunoprecipitation experiment in presence of the drugs (Figure 3A,B).

**Figure 3:**
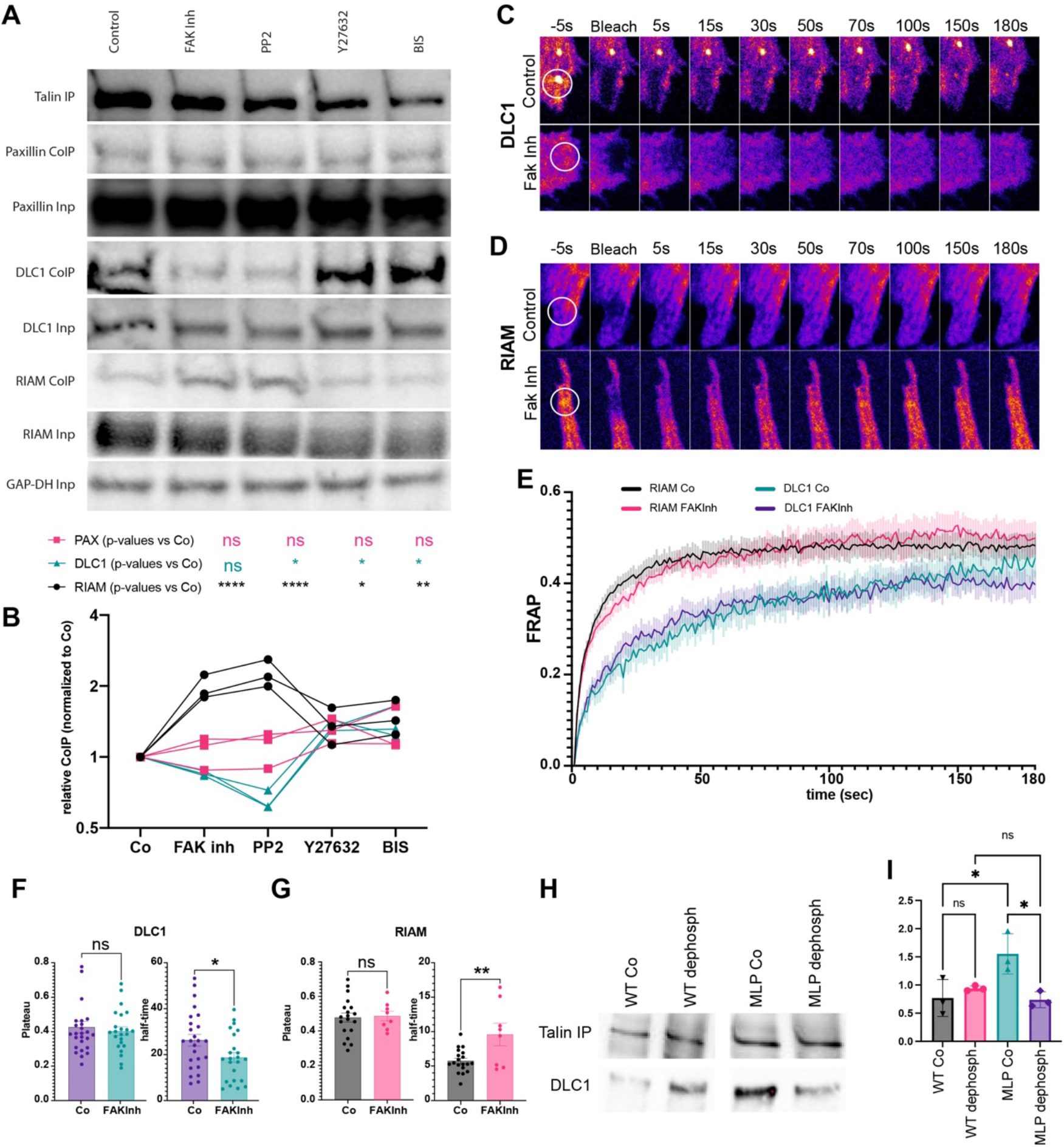
The talin interactome is regulated through phosphorylation. **A)** Co-immunoprecipitation analysis for talin interactions. Neonatal rat cardiomyocytes were cultured on PDMS with 6kPa stiffness for 7 days, treatment with the indicated inhibitors and immunoprecipitation with an anti-talin antibody in presence of the indicated inhibitors. Samples were then subjected to western blotting and probed with antibodies against paxillin, RIAM and DLC1, indicating changes in the talin interactions after FAK and Src inhibition. **B**) quantification from three independent repeats; **C-G)** Fluorescence recovery after photobleaching (FRAP) assays of DLC1 (C) and RIAM (D) in neonatal rat cardiomyocytes. **E)** average recovery curves (mean +SEM); **F)** Quantification of plateau (mobile fraction) and recovery half-time for DLC1 and **G)** RIAM. **H**,**I)** Immunoprecipitations from WT and MLP knockout mouse hearts, with or without treatment with alkaline phosphatase indicates disease and phosphorylation dependent changes also in vivo. *p<0.0332, **p<0.0021, ***p<0.0002, ****p<0.0001; p-values from one-way ANOVA with Dunnett correction for multiple comparisons (B,I) or unpaired t-test (F,G);

Interestingly, Src and FAK inhibition altered the talin interactions, whereby the bound fraction of RIAM was increased and the interaction with DLC1 was simultaneously reduced, suggesting that the competition between DLC1 and RIAM for the talin binding site was dependent on integrin related signalling through Src and FAK. Noteworthy, we did not find any changes to paxillin binding and the inhibition of Rho Kinase and PKC showed only minor effects (both slightly increasing the binding of DLC1 and RIAM, compared to the control). Again, we turned to FRAP experiments to observe the changes of DLC1 and RIAM adhesion binding after FAK and Src inhibition *in cellulo* (Fig 3C-G). Indeed, the changes to the protein dynamics agreed with the results from the co-immunoprecipitations, suggesting an increased half-time of recovery for RIAM and a reduced half-time for DLC1 after FAK inhibition (Fig 3F,G). Again, the mobile fraction was unchanged, suggesting that the phosphorylation alters the interaction lifetime through altering the affinities and competition for binding, rather than completely inhibiting the binding of DLC1 or RIAM. Western blotting for active phosphorylated FAK further indicated that FAK activity was highest on a healthy adult heart stiffness (6kPa; Supplementary Fig S3). Due to the disruptive effect of Src inhibition on the cardiomyocyte adhesions morphology, we were not able to reliably evaluate the changes after Src inhibition.

Since our data showed that DLC1 expression was increased in the MLP knockout mouse (Fig. 1D), with simultaneous increase in RIAM (Fig. S2A,D), we wanted to test if this was reflected also in altered DLC1 binding to talin. For this we performed co-immunoprecipitation assays from wild type and MLP knock out hearts at 6 month of age, where increased stiffness and fibrosis are evident^39^. This experiment showed an increase in DLC1 talin interaction in the fibrotic MLP knock out heart (Fig 3H,I). Importantly, due to a lower healthy heart stiffness in mice (∼ 1.5kPa diastolic and 3kPa systolic stiffness in mice, compared to 15.8 ± 2.8 kPa systolic stiffness in human, as previously measured by shear wave elastometry^40, 41^), this result was consistent with the changes we observed in our immuno-precipitations from the NRCs on different stiffness, where the DLC1-talin interaction was lower at 1kPa, compared to 6kPa.

To test if the increased talin binding might be imprinted through altered phosphorylation also *in vivo* we treated a part of the samples with alkaline phosphatase which removes tyrosine and to a lesser degree also serine/threonine phosphorylation^42^. Indeed, the immunoprecipitations revealed that the increased binding of DLC1 in MLP knockout samples was reduced after dephosphorylation, confirming that DLC1 binding to talin was regulated through phosphorylation also *in vivo*.

### DLC1 and RIAM are directly competing for the binding to R8 in cardiomyocytes

RIAM can bind both the R3 and R8 domain but the FRAP experiments indicated a competition in the binding for R8, with opposite trends for RIAM and DLC binding through increased ECM stiffness and FAK inhibition. To test directly for the competition and its alteration through phosphorylation, we decided for an optogenetic approach. The LOVTRAP system uses a small protein tag (Zdark, or Zdk) that has been engineered to selectively bind the LOV2 photosensor domain from *Avena sativa* phototropin 1 in the dark state only. The LOV2 domain is linked to a fragment of the TOM20 mitochondrial anchoring sequence^32^ and hence sequesters Zdk-tagged proteins to the mitochondria in the dark state. Upon illumination with blue light the Zdk-tagged protein is rapidly released from the mitochondria and can interact with its typical interaction partners. This system is therefore ideally suited to investigate a direct competition between RIAM and DLC1 at the R8 domain (Fig 4A). Because of the technically challenging nature, we performed the experiment in the C2C12 muscle cell line and on glass coverslips. As expected, when cells were transfected with a combination of DLC1-GFP, RIAM-mCherry-ZDK and TOM20-LOV and cells were kept under dark conditions, RIAM was localised to mitochondria. Illumination with blue light immediately dislocated RIAM from the mitochondria and resulted in an increased adhesion localisation (Fig 4B,C). Quantification of RIAM and DLC1 suggested a simultaneous decrease in the amount of adhesion localised DLC1, which was peaking 3s after illumination, confirming a competition of RIAM with DLC1 for adhesion localisation. Intriguingly, DLC1 quickly returned to pre-stimulation levels (6s). Treatment with FAK inhibitor did not affect the adhesion localisation per se, but resulted in a change in the dynamics, especially a prolongated enrichment of RIAM at the adhesions and delayed return of displaced DLC1 to the adhesion, overall confirming a shift in the affinities towards stronger RIAM and reduced DLC1 binding to talin (Fig 4B-H).

**Figure 4:**
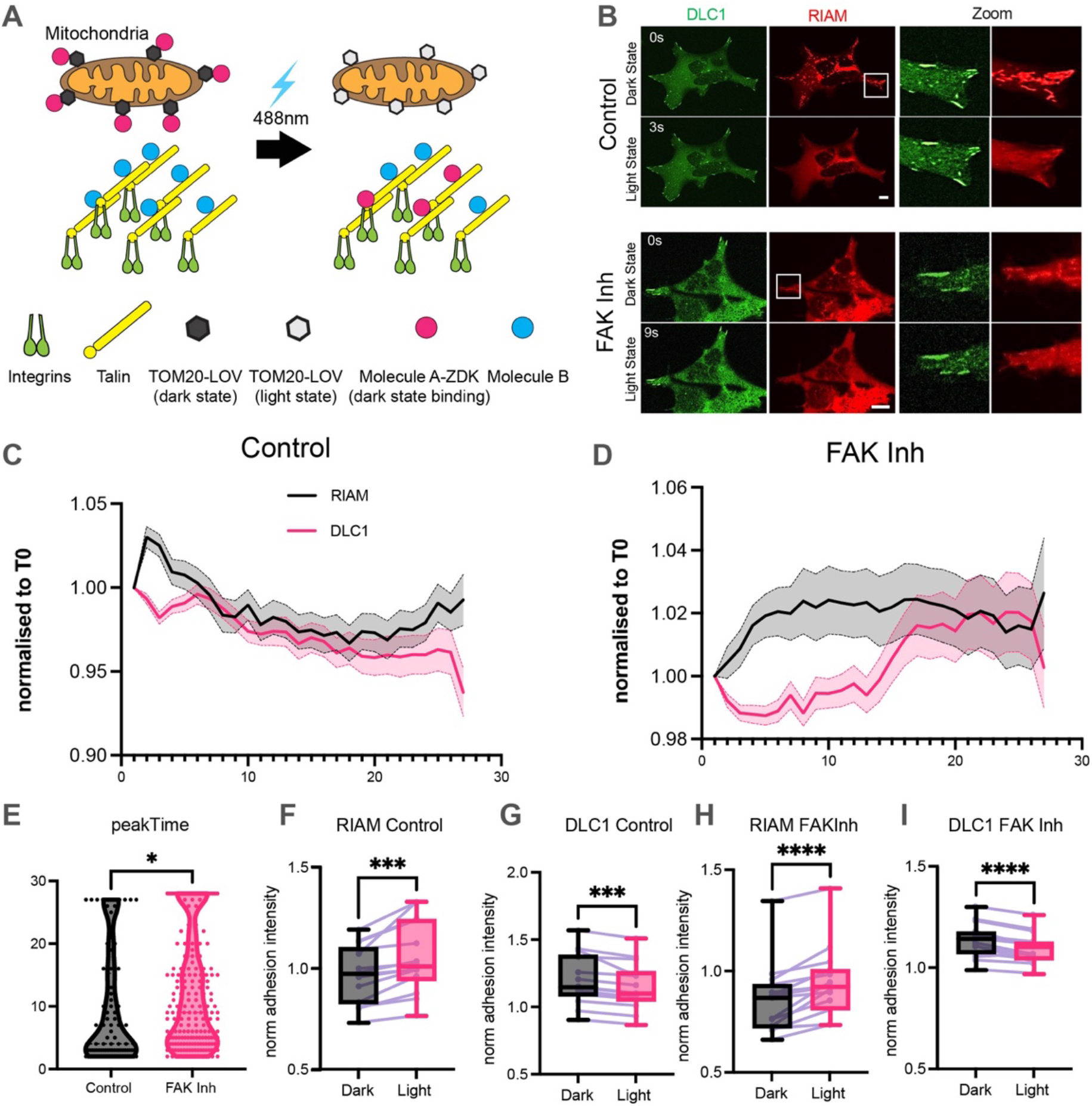
DLC1 and RIAM are directly competing for talin binding. **A)** Schematic of the LOVTRAP system. Illumination with green light converts the LOV domain into the light state and displaces the dark-state binding ZDK tag and associated molecule (here RIAM) from the mitochondria to enable adhesion binding. B) Imaging of cells with a green laser immediately displaces RIAM from a mitochondria localisation, leading to increased adhesion binding of RIAM and reduced adhesion binding of DLC1 (quantified in (C)). The dynamics is changed after FAK inhibition (quantified in (D)) and also apparent form the change in the time until the peak in RIAM binding to the adhesion is reached (E). F-I) Adhesion intensities (normalised to whole cell intensities) at the time of peak adhesion enrichment for RIAM, displayed as box plot and changes for each cell from 3 independent experiments. *p<0.0332, **p<0.0021, ***p<0.0002, ****p<0.0001; p-values from paired t-test;

### Loss of DLC1 leads to stiffness dependent changes in RhoA activity and sarcomeric structure

Because DLC1 appears to be the dominant RhoGAP in cardiomyocytes, we next wanted to test if it was involved in regulating RhoA activity in cardiomyocytes. For this, we knocked down DLC1 in NRCs using siRNA (Fig 5A) and measured the RhoA activity by fluorescent lifetime imaging of the RhoA biosensor. This single chain biosensor is based on the ability of the Rho-binding domain to interact with active GTP-RhoA, but not with inactive GDP-RhoA. The interaction brings the fluorophores (CFP as FRET donor and YFP as FRET acceptor) in close proximity and thus results in a shortening of the fluorescent lifetime of the donor in the active GTP bound state (Fig 5B)^43^. Consistent with a higher amount of DLC1-talin interaction on 6kPa, we found a longer lifetime, i.e. reduced activity of RhoA in cardiomyocytes cultured on 6kPa, compared to 130kPa. Moreover, knock down of DLC1 increased the RhoA activity (i.e. decreased the lifetime) on 6kPa to levels comparable to what was detected on 130kPa, while the knockdown on 130kPa had no additional effect on the lifetime (Fig 5C-E). When we assessed the effect of the knock down on myofibrillar structures, using α-actinin as a marker for sarcomeric structures, we further found a reduced number of cells displaying fully mature sarcomeres on 6kPa, and an increase of cells displaying myofibrillar disarray (Fig 5F-G).

**Figure 5:**
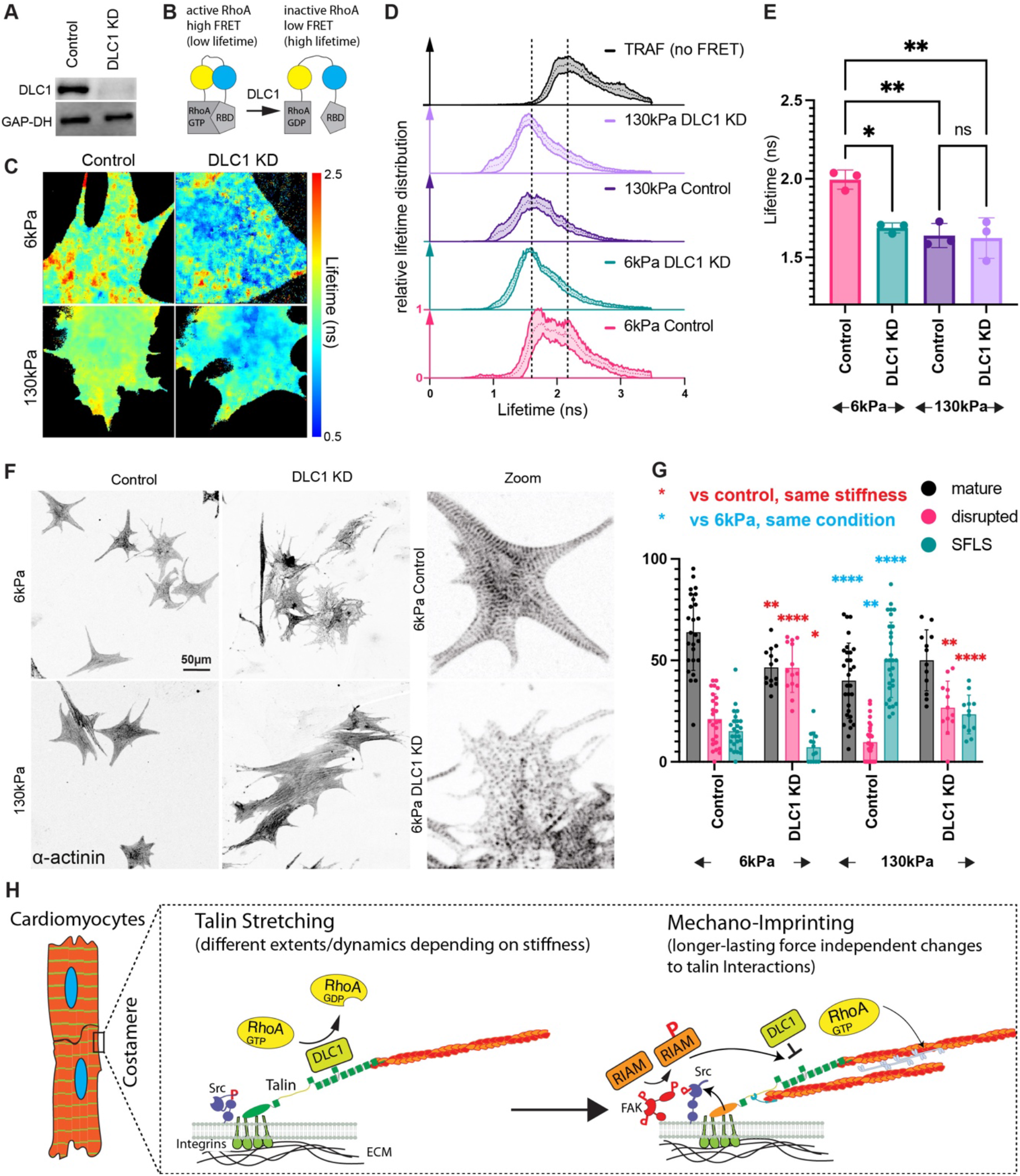
Loss of DLC1 leads to a stiffness dependent reduction in RhoA activity and sarcomeric organisation. **A)** DLC1 siRNA validation. **B)** Schematic for RhoA biosensor, adapted from ^43^. **C)** Example fluorescence lifetime images from 3 independent repeats; **D)** distribution of lifetimes per pixel; **E)** peak lifetime after log-normal fit, averaged per repeat; **F)** a-actinin staining in neonatal rat cardiomyocytes indicates a loss of sarcomeres after DLC1 knockdown, quantified in **(G). H)** Schematic. Different integrin signalling leads to different levels of Src and FAK activation and phosphorylation of RIAM and/or DLC1. These posttranslational modifications lead to longer-lasting, imprinted changes to the affinities and dynamicity of binding to the talin R8 domain and thus, modify the level of RhoA activity. Because of the importance of DLC1 for fine-tuning the levels of RhoA activity, this impacts the cardiomyocyte maturity and disease progression. *p<0.0332, **p<0.0021, ***p<0.0002, ****p<0.0001; p-values from one-way ANOVA with Dunnett correction for multiple comparisons (E) or two-way ANOVA with Tukey correction for multiple comparisons (G);

Cardiomyocytes on 130kPa displayed overall a larger proportion of cells containing long stress-fibre like structures even in the control condition (as previously reported)^17^ which however was shifted towards disrupted sarcomeres after DLC1 knock down.

Since our data indicated that under certain conditions (MLP knock out heart), DLC1 binding to talin was increased we further tested for the effect of DLC1 overexpression (Fig. S4). Intriguingly, adenoviral driven overexpression also reduced the maturity of the cardiomyocytes, but in this case led to an increase in the number of cells with excessive stress-fibre like structures, reaching a level that was otherwise seen on fibrotic stiffness (130kPa, with and without overexpression of DLC1). Overall, this suggests that mechanical memory through the talin interactome leads to alterations in DLC1 activity and further the level of DLC1 at adhesions finely tunes the activity of RhoA in a stiffness dependent way to affect sarcomeric integrity, cardiomyocyte function and pathological cardiac remodelling^26, 31^.

## Discussion

While mechanical signalling has been well described as critical factor influencing health and disease ^44^, the question how mechanical signals can be memorized by cells or tissues is less clear. Here our data shows a novel mechanism of mechanical memory through a modification of talin interactions. We find that DLC1 is the key RhoGAP in cardiomyocyte adhesions and interacts with talin in a stiffness dependent way. This interaction is regulated through changing affinities of RIAM and DLC1 for the shared binding site. Importantly these changes are imprinted and remain in place even in absence of tension on talin. Moreover, we confirm this mechanical memory in vivo, in fibrotic mouse hearts from the MLP knockout model for dilated cardiomyopathy.

Only recently, research began to elucidate the existence and regulation of mechanical memory. Several studies indicated that stiffness was not only sensed instantaneously, but affected the behaviour even after subsequent change of stiffness and as such demonstrated mechanical memory^45-50^. Mesenchymal stem cells were found to remember the previous exposure to stiff surfaces, even when the stiffness was reduced^45^. Similarly, culturing lung fibroblasts on stiff substrates primed the cells for sustained myofibroblast activity, even after returning these onto soft surfaces for two weeks^46^. Also, oral squamous cell carcinoma cells were primed by a stiff surface and remained migratory even when their environment softened^50^. On the other hand, adipose-derived stem cells could be primed by culturing on a soft surface, which then led to a delayed response to increased stiffness^47^. In addition to YAP/TAZ signalling, FAK signalling and epigenetic modifications have been suggested as a mechanism for storing the information^45, 48-50^.

Even though there is growing evidence for mechanical memory, its regulation in cardiomyocytes has so far not received much attention. However, there is ample evidence for the interplay of mechanical information and cardiac disease, making it likely that mechanical memory could be at play in cardiomyocytes, as well. Indeed, a previous study of heart failure found that after anti-fibrotic treatment, cardiomyocyte hypertrophy persisted^51^. This suggests an uncoupling of mechanical signals from mechanotransduction, and points to the existence of mechanical memory at cellular and/or molecular level. In general, cardiomyocytes sense the mechanical properties of their environment and in response adapt their behaviour and function^1, 17^. Especially, cardiac maladaptation and disease progression can occur through disruption of mechanosensing through changes to: i) contractile force, ii) stiffness or iii) adhesion composition; i) Contractile forces are modified through the altered expression of motor proteins and their regulators as well as the changing load and its effect on the cardiac output ^1, 52, 53^, which can lead to cardiac remodelling and to heart failure after systolic dysfunction^54^. ii) Altered passive stiffness can lead to diastolic dysfunction ^55^. Here, especially the transition from the compensated stage to diastolic heart failure is linked with progressive myocardial stiffening^56-58^ and has been associated with collagen crosslinking, through lysyl oxidases (LOXs) and LOX like enzymes (LOXLs) ^1, 6-9, 51, 59^. iii) Cardiomyocytes connect to the extracellular matrix (ECM) via integrins at costameres, however integrin expression is modulated as an adaptive response to heart disease^1, 60-67^. Changing integrin isoforms affect adhesion formation, force transduction and binding of other adhesion proteins, such as the mechanosensitive protein talin^1, 10^. Especially talin is a as an integral component of the cardiac costamere (as of focal adhesions) and a key protein in the heart^68-70^. Talin is required for cardiomyocyte remodelling during heart growth^70^. Moreover, loss of talin leads to costameric instability and cardiomyopathy^68^.

Importantly, both talin isoforms modify the hypertrophic response^69^. Indeed, talin was previously proposed to be ideally placed as a hub also for mechanical memory^37^,^71^. In the MeshCODE theory, each talin helix bundle is proposed to act as a binary switch which can store mechanical information in the talin molecule^71^. Changes in tension, as previously observed in cardiomyocytes would switch individual domains, lead to alteration in the force dependent binding and alter signalling^71^. This proposed mechanism is indeed in agreement with our findings, albeit our data suggests an additional regulation through posttranslational modifications, especially phosphorylation through FAK and Src, leading to changes in the affinities and competition for talin binding between DLC1 and RIAM. Talin binding to integrin and the actin cytoskeleton results in the forces for the talin stretching, further also triggering the outside-in signalling cascade that includes the activation of kinases including FAK and Src^72, 73^. A recent study found that mechanical memory depended on FAK activity in oral squamous carcinoma cells^50^, and indeed DLC1 activity depends on a motif that forms a complex with talin and also FAK^74^. Moreover, FAK and Src phosphorylation are known to release autoinhibition of RIAM allowing RAP1 binding and plasma membrane association^75^. Here increased binding of RIAM to talin R8 domain after FAK and Src inhibition might relate to a decrease in the plasma membrane association, since the R8 domain will likely be extended away from the membrane according to studies of focal adhesion 3D nano structure^75, 76^. Thus, in our context, FAK inhibition caused a decrease in DLC1 binding to talin and increased RIAM binding in co-immunoprecipitations and higher dynamics in FRAP experiments. Matching with this, western blotting further showed elevated levels of active pFAK in NRCs at 6kPa, consistent with higher levels of DLC1-talin binding.

The direct competition is consequently depending on FAK phosphorylation as also evidenced by our optogenetic LOVTRAP approach. Overall this has a strong impact on downstream signalling. DLC1 knock down increases RhoA activity and at the same time leads to myofibrillar disruptions, while on the other hand an increase in DLC1 expression is linked with the formation of long stress-fibre like structures. This suggests that RhoA activity in cardiomyocytes needs to be finely tuned and indeed the literature points to both benefits and detrimental effects of RhoA in cardiovascular health and disease ^31^. RhoA is upregulated in early heart development^29^, but RhoA overexpression in the adult heart results in dilated cardiomyopathy (albeit only at high, not at moderate over-expression)^30, 77^. Cardiomyocyte specific knock out of RhoA on the other hand, resulted in an accelerated and more severe dilated cardiomyopathy after chronic pressure overload^26^.

Importantly, the pathway we outline here is likely only partially reflecting the complexity of the talin mechanical interactions and memory, whereby posttranslational modifications affect force transduction and sensing. For instance, methylation of R13 by Ezh2 has been shown to disrupt actin binding^78^ and hence force transduction. On the other hand, cyclin dependent kinase 1 (CDK1) was described to bind to the R8 domain and CDK1 phosphorylation of a serine residue (S1589) in the linker between R7 and R8 reduces the force needed for unfolding of the R8 domain^79^.

Overall our results show that mechanical sensing and activation of integrin related kinase pathways, including FAK and Src, alter the talin interactome and through this imprint the mechanical information, to determine RhoA activity levels and can influence the cardiomyocyte cytoskeleton, function and cardiac remodelling in heart disease.

## Methods

### Antibodies and Reagents

The following primary antibodies were used in the study: Talin (Abcam ab71333), alpha-actinin (Sigma, A7811), Vinculin (Sigma, V9131), DLC1 (NBP1-88824PEP, NovusBio), paxillin (BD Biosciences, 610052), RIAM (Abcam, ab92537) GAPDH (Abcam, ab8245).

The following reagents were used at the listed concentrations: Bisindolylmaleimide I (BIS I, Sigma, used at 500nM), PP2 (Cayman Chemical, 10 μM), Y27632 (Enzo Life Sciences, 10 μM), FAK Inhibitor 14 (Cayman Chemical, 1 µg/ml).

pTriEx-NTOM20-LOV2 (Addgene plasmid # 81009) and pTriEx-mCherry-Zdk1 (Addgene plasmid # 81057) were a gift from Klaus Hahn^32^. The RhoA-FRET biosensor was a gift from O. Pertz^43^.

Adenoviral constructs for DLC1-GFP (based on the sequence NM_001348081.2), RIAM-GFP (based on the sequence NM_019043.4), and RIAM-mChe-Zdk1 (based on pTriEx-mCherry-Zdk1 and NM_019043.4) were generated by Vectorbuilder.

### Human tissues

Surplus human tissue from patients undergoing surgery was snap frozen in liquid nitrogen. For patient characteristics see Suppl. Table 1. Samples from left ventricular assist device (LVAD) implantation and explanted hearts (HTx) were in end stage heart failure (ejection fraction, EF, < 40 %), while aortic stenosis (AoS) samples (apart from one exception) had preserved cardiac function (EF > 50 %).

### Mouse tissues

MLP knock out mice, a genetic model for heart failure, and wild type littermates were sacrificed at 6 months of age and hearts were snap frozen for cryosections or protein extraction.

### PDMS Substrates

Polydimethylsiloxane (PDMS) substrates were prepared, by mixing Sylgard 527 with Sylgard 184 as described previously^80^. The stiffness of the mixtures was measured by rheology as described previously^*17*^. After spin-coating onto coverslips, the PDMS was cured either at 80°C for 2h or 60°C overnight.

### Cardiomyocyte isolation, culture and siRNA

Neonatal Rats Cardiomyocytes were isolated using the sequential digestion method as described previously^81^. Briefly, hearts were dissected into ice-cold ADS buffer (116 mM NaCl, 20 mM Hepes, 0.8 mM NaH2PO4, 5.6 mM glucose, 5.4 mM KCL, 0.8 mM MgSO4). After hearts settled down to the bottom, they were washed once with ADS buffer. ADS buffer was then removed, and hearts were incubated with 5 ml enzyme solution in ADS (ES, 246U collagenase and 0.6 mg/ml pancreatin), for 5 min, at 37°C under vigorous shaking. Supernatant was discarded. This step was followed by five to six digests, until hearts were completely digested. Each time, 5 ml fresh enzyme solution (ES) was added to the hearts and incubated 15 min at 37°C, under shaking. Hearts were pipetted up and down 30 times using a pasteur pipette. After settling down, supernatant was transferred into plating medium (65% DMEM, 17% M199, 10% horse serum, 5% fetal bovine serum (FBS), 2% GlutaMAX, 1% penecillin/streptamycin (P/S)). Two digests each were combined in one tube with 20 ml plating medium, then cleared through a 100 μm cell strainer and spun down at 1200 r.p.m. for 5 min at RT, before being resuspended in 10 ml plating medium. Cells were pooled together and pre-plated for 90 min to enrich the cardiomyocytes. Cardiomyocytes were then plated onto the respective substrates as indicated in the text or figures. Medium was changed the next day to maintenance medium (77% DMEM, 18% M199, 2% horse serum, 2% GlutaMAX, 1% P/S), or serum starvation medium (as above, but excluding the horse serum).

DLC1 knockdown was performed by using 1µM DLC1 Accell siRNA SMART Pool (mix of 4 siRNA targeting the DLC1 sequence GUAUGAGCAUCUAUGACAA, CUCCUGUCCAGAUAUUUGA, CUAUUAUACCUGGUUUAGC and GUUUUAGCAUGAAAGGUCA) #E-088843-01-0010 from Horizon, which requires no transfection reagent for delivery. The scrambled siRNA was a non-targeting siRNA Accell #D-001910-01-05. Cells were incubated for 48 h after transfection.

### Immunofluorescence

Hearts were covered with optimal cutting temperature (OCT) for cryosectioning on a cryostat (Leica Biosystems). Frozen sections (12 μm thick) were fixed in ice-cold acetone at -20°C for 5min before rehydration in PBS and blocking for 30 min with 5% goat serum in PBS. Neonatal rat cardiomyocytes (NRCs) were fixed by using 4% paraformaldehyde (PFA) for 10min at room temperature, permeabilized with 0.2% Triton X-100 for 5min at room temperature, and blocked for 1h at room temperature with 5% goat serum. Immunostaining of hearts sections and NRCs on PDMS were performed by incubating the samples with primary antibodies at 4°C overnight in a wet chamber. On the next day, after three 5min washings with PBS, samples were incubated for 1hour with secondary antibodies, Phalloidin and DAPI at room temperature. Samples were washed three times before mounting using Fluoromount Mounting Media (Thermo Fisher Scientific).

### Microscopy

Optogenetic and FRAP experiments were performed a Nikon CSU-W1 SoRA Spining Disk confocal microscope with two Photometrics Prime BSI cameras and equipped with environmental chamber for CO2 and temperature control, using a 60× oil Lambda Apochromat objective and after transfection of the NRCs for 48h. For FRAP, 5 frames at 1fps were acquired to record the baseline, before laser stimulation of 3µm circular ROIs on focal adhesion sites at 10% laser power for 200msec. Recovery after photobleaching was observed for 3min with acquisition every second. For optogenetic experiments, cells were located and then left 3min in dark to restore the mitochondrial localisation of ZDK-mCherry-RIAM. Images were acquired for 30s at 1fps using both 488 and 568nm laser lines, whereby the first 488nm exposure was sufficient to convert LOV into the light state and displace ZDK-mCherry-RIAM from the mitochondria.

FRET acquisitions of the single chain RhoA-FRET biosensor were performed on a custom built multi-beam confocal FLIM system as described previously^82^.

### Western blotting and co-immunoprecipitation

Snap frozen cardiac samples (either from human patients at time of surgery or from mice culled by cervical dislocation, after quick rinse in PBS) were powdered with a liquid nitrogen cooled BioPulverizer100 (BioSpecProducts). Samples were mixed with 2xSDS sample buffer (125 mM Tris HCpH 6.8, 200 mM DTT, 4 % SDS, 12 % glycerol, 0.05 % bromophenolblue, using 200 uL of 2xSDS sample buffer per 100 ug tissue), heated at 65 degrees for 15 min and then sonicated and subjected to western blotting.

Neonatal rat cardiomyocyte protein lysates were extracted using RIPA Buffer with phosphatase and protease inhibitors. Lysate for co-immunoprecipitation (Co-IP) were extracted using and Pierce IP Lysis Buffer (87788). Co-IPs were performed by using the magnetic Sure Beads Protein A from BioRad washed with dilution buffer (10mM Tris, 150mM NaCl, 0.5mM EDTA PH=7.4, and coated with the Talin antibody (Abcam ab71333 5µg/200mL). The protein lysates were added to the beads for 1h at room temperature before being washed with washing buffer (10mM Tris, 150mM NaCl, 0.5mM EDTA, 0.05% Triton x100) and eluted in glycine 20mM PH =2 for 5min at room temperature and neutralized with 1M Tris, PH=10.4. The protein dephosphorylation was performed on total WT and MLP knockout heart lysates by using Alkaline Phosphatase (rSAP)(M0371S NEB, 1U of rSAP/10mg of protein). The protein lysate containing phosphatases, but omitting phosphatase inhibitors were incubated 30min at 37°C, before subjecting the samples to the co-immunoprecipitation as described above.

Samples were separated on Mini-PROTEAN TGX Precast Gels (4-15%, Biorad), loading was adjusted based on intensity of bands after Coomassie stain. For western blotting, samples were separated in gels and transferred onto nitrocellulose, or PVDF membranes. Transfer was verified by PonceauS stain (0.1% Ponceau, 0.5% acetic acid, MERCK). The membranes were cut depending on the molecular weights of the desired proteins and rinsed in 0.1% TBS tween. A 5% milk in TBS tween blocking buffer was then applied to the membrane for 1 hour at room temperature. Primary antibodies were incubated at 4ºC overnight or 1h at room temperature for GAPDH and HRP-conjugated secondary antibodies were incubated 1h at room temperature.

### Quantification and Statistical analysis

Quantification were performed using ImageJ and Matlab. ROI were drawn around the area of laser stimulation for FRAP experiments, as well as in a reference location with no bleaching and another one for background. The background intensity was subtracted from the ROI intensity of the photobleaching spot and divided by the reference intensity. The 5 seconds preceding laser stimulation were averaged, and data was normalized by subtracting the value of the laser stimulation at T0 from every time-points and divided by the average of the 5sec before stimulation. For quantification of the optogenetic experiments, focal adhesion sites were precisely selected to quantify the intensity after light stimulation. Each value of the time series have been divided by the intensity value of the whole cell. Western Blots were quantify using Image Lab (Biorad) or ImageJ and normalized to GAPDH expression. Protein expression in talin Co-IP was normalized on talin expression. The maturity of sarcomeres was quantified using α-actinin expression based on sarcomere morphology and visual presence of stress fiber-like structures or disrupted staining pattern (missing sarcomeres, dot like z-discs). All statistical tests are indicated in the figure legends. Data sets were tested for normal distribution using the Shapiro–Wilk test. All statistical tests were performed with GraphPad prism.

## Ethics Statement

Animal studies have been performed in accordance with the ethical standards laid down in the 1964 Declaration of Helsinki and its later amendments. Experimental procedures were performed in accordance with the Directive 2010/63/EU and UK Home office guidelines (project licences P572C7345 and PDCE16CB0) and approved by the respective institutional ethical review boards.

Ethics for collection of human tissue during surgery for the purposes of establishment of a biobank for left-over biological materials and associated data and experimental use were obtained at the UOC di Cardiochirurgia Verona – Protocol code: BB-CCH, Proj. 847CESC.

## Author Contributions

Conceptualization, T.I.; Methodology, E.M., I.X., J.L. and T.I; Investigation: E.M., I.X., P.P., A.A., M.R. and T.I.; Writing – Original Draft, Review & Editing, E.M., K.G. and T.I.; Resources, K.G., D.P. S.A.B., E.E. and T.I.; Supervision, K.G., S.A.B. and T.I.

## Acknowledgements

We would like to thank Ben Goult (University of Kent) and Sanjay Sinha (University of Cambridge) for helpful discussions and comments on the text and figures. We further thank Klaus Hahn (University of North Carolina at Chapel Hill) for discussions related to the LOVTRAP construct.

## References

1. Ward, M. & Iskratsch, T. Mix and (mis-)match - The mechanosensing machinery in the changing environment of the developing, healthy adult and diseased heart. Biochim Biophys Acta Mol Cell Res 1867, 118436 (2020).

2. Yamauchi, M. & Sricholpech, M. Lysine post-translational modifications of collagen. Essays Biochem 52, 113–133 (2012).

3. Ruotsalainen, H., Sipila, L., Kerkela, E., Pospiech, H. & Myllyla, R. Characterization of cDNAs for mouse lysyl hydroxylase 1, 2 and 3, their phylogenetic analysis and tissue-specific expression in the mouse. Matrix Biol 18, 325–329 (1999).

4. Lopez, B. et al. Role of lysyl oxidase in myocardial fibrosis: from basic science to clinical aspects. Am J Physiol Heart Circ Physiol 299, H1–9 (2010).

5. Maki, J.M. Lysyl oxidases in mammalian development and certain pathological conditions. Histol Histopathol 24, 651–660 (2009).

6. Hermida, N. et al. A synthetic peptide from transforming growth factor-beta1 type III receptor prevents myocardial fibrosis in spontaneously hypertensive rats. Cardiovasc Res 81, 601–609 (2009).

7. Hughes, W.M., Jr. et al. Role of copper and homocysteine in pressure overload heart failure. Cardiovasc Toxicol 8, 137–144 (2008).

8. McCormick, R.J., Musch, T.I., Bergman, B.C. & Thomas, D.P. Regional differences in LV collagen accumulation and mature cross-linking after myocardial infarction in rats. Am J Physiol 266, H354–359 (1994).

9. Lopez, B. et al. Impact of treatment on myocardial lysyl oxidase expression and collagen cross-linking in patients with heart failure. Hypertension 53, 236–242 (2009).

10. Sit, B., Gutmann, D. & Iskratsch, T. Costameres, dense plaques and podosomes: the cell matrix adhesions in cardiovascular mechanosensing. J Muscle Res Cell Motil 40, 197–209 (2019).

11. Ervasti, J.M. Costameres: the Achilles’ heel of Herculean muscle. J Biol Chem 278, 13591–13594 (2003).

12. Engler, A.J. et al. Embryonic cardiomyocytes beat best on a matrix with heart-like elasticity: scar-like rigidity inhibits beating. Journal of cell science 121, 3794–3802 (2008).

13. McCain, M.L., Yuan, H., Pasqualini, F.S., Campbell, P.H. & Parker, K.K. Matrix elasticity regulates the optimal cardiac myocyte shape for contractility. Am J Physiol Heart Circ Physiol 306, H1525–1539 (2014).

14. Hazeltine, L.B. et al. Effects of substrate mechanics on contractility of cardiomyocytes generated from human pluripotent stem cells. Int J Cell Biol 2012, 508294 (2012).

15. Hersch, N. et al. The constant beat: cardiomyocytes adapt their forces by equal contraction upon environmental stiffening. Biol Open 2, 351–361 (2013).

16. Rodriguez, A.G., Han, S.J., Regnier, M. & Sniadecki, N.J. Substrate stiffness increases twitch power of neonatal cardiomyocytes in correlation with changes in myofibril structure and intracellular calcium. Biophys J 101, 2455–2464 (2011).

17. Pandey, P. et al. Cardiomyocytes Sense Matrix Rigidity through a Combination of Muscle and Non-muscle Myosin Contractions. Dev Cell 44, 326–336 e323 (2018).

18. Goult, B.T., Brown, N.H. & Schwartz, M.A. Talin in mechanotransduction and mechanomemory at a glance. Journal of Cell Science 134 (2021).

19. Klapholz, B. & Brown, N.H. Talin - the master of integrin adhesions. J Cell Sci 130, 2435–2446 (2017).

20. Yao, M. et al. The mechanical response of talin. Nat Commun 7, 11966 (2016).

21. del Rio, A. et al. Stretching single talin rod molecules activates vinculin binding. Science 323, 638–641 (2009).

22. Goult, B.T. et al. RIAM and vinculin binding to talin are mutually exclusive and regulate adhesion assembly and turnover. J Biol Chem 288, 8238–8249 (2013).

23. Haining, A.W.M. et al. Mechanotransduction in talin through the interaction of the R8 domain with DLC1. PLoS Biol 16, e2005599 (2018).

24. Kaushik, S., Ravi, A., Hameed, F.M. & Low, B.C. Concerted modulation of paxillin dynamics at focal adhesions by deleted in liver cancer-1 and focal adhesion kinase during early cell spreading. Cytoskeleton 71, 677–694 (2014).

25. Erlmann, P. et al. DLC1 Activation Requires Lipid Interaction through a Polybasic Region Preceding the RhoGAP Domain. Molecular Biology of the Cell 20, 4400–4411 (2009).

26. Lauriol, J. et al. RhoA signaling in cardiomyocytes protects against stress-induced heart failure but facilitates cardiac fibrosis. Sci Signal 7, ra100 (2014).

27. Torsoni, A.S., Fonseca, P.M., Crosara-Alberto, D.P. & Franchini, K.G. Early activation of p160ROCK by pressure overload in rat heart. Am J Physiol Cell Physiol 284, C1411–1419 (2003).

28. Phrommintikul, A. et al. Effects of a Rho kinase inhibitor on pressure overload induced cardiac hypertrophy and associated diastolic dysfunction. Am J Physiol Heart Circ Physiol 294, H1804–1814 (2008).

29. Kaarbø, M., Crane, D.I. & Murrell, W.G. RhoA is highly up-regulated in the process of early heart development of the chick and important for normal embryogenesis. Developmental dynamics: an official publication of the American Association of Anatomists 227, 35–47 (2003).

30. Xiang, S.Y. et al. RhoA protects the mouse heart against ischemia/reperfusion injury. J Clin Invest 121, 3269–3276 (2011).

31. Kilian, L.S., Voran, J., Frank, D. & Rangrez, A.Y. RhoA: a dubious molecule in cardiac pathophysiology. Journal of Biomedical Science 28, 33 (2021).

32. Wang, H. et al. LOVTRAP: an optogenetic system for photoinduced protein dissociation. Nat Methods 13, 755–758 (2016).

33. Muller, P.M. et al. Systems analysis of RhoGEF and RhoGAP regulatory proteins reveals spatially organized RAC1 signalling from integrin adhesions. Nat Cell Biol 22, 498–511 (2020).

34. Goult, B.T., Yan, J. & Schwartz, M.A. Talin as a mechanosensitive signaling hub. Journal of Cell Biology 217, 3776–3784 (2018).

35. Hirth, S. et al. Paxillin and Focal Adhesion Kinase (FAK) Regulate Cardiac Contractility in the Zebrafish Heart. PLOS ONE 11, e0150323 (2016).

36. Gehmlich, K. et al. Paxillin and ponsin interact in nascent costameres of muscle cells. J Mol Biol 369, 665–682 (2007).

37. Goult, B.T., Brown, N.H. & Schwartz, M.A. Talin in mechanotransduction and mechanomemory at a glance. J Cell Sci 134 (2021).

38. Tripathi, B.K. et al. SRC and ERK cooperatively phosphorylate DLC1 and attenuate its Rho-GAP and tumor suppressor functions. Journal of Cell Biology 218, 3060–3076 (2019).

39. Unsöld, B. et al. Age-dependent changes in contractile function and passive elastic properties of myocardium from mice lacking muscle LIM protein (MLP). European Journal of Heart Failure 14, 430–437 (2012).

40. Liu, Y., Royston, T.J., Klatt, D. & Lewandowski, E.D. Cardiac MR elastography of the mouse: Initial results. Magn Reson Med 76, 1879–1886 (2016).

41. Kolipaka, A. et al. in Proceedings of the 19th Annual Meeting of ISMRM, Montreal, Canada 274 (2011).

42. Takahashi, K. et al. Tyrosine-specific dephosphorylation-phosphorylation with alkaline phosphatases and epidermal growth factor receptor kinase as evidenced by 31P NMR spectroscopy. J Biochem 101, 1107–1114 (1987).

43. Pertz, O., Hodgson, L., Klemke, R.L. & Hahn, K.M. Spatiotemporal dynamics of RhoA activity in migrating cells. Nature 440, 1069–1072 (2006).

44. Iskratsch, T., Wolfenson, H. & Sheetz, M.P. Appreciating force and shape-the rise of mechanotransduction in cell biology. Nat Rev Mol Cell Biol 15, 825–833 (2014).

45. Yang, C., Tibbitt, M.W., Basta, L. & Anseth, K.S. Mechanical memory and dosing influence stem cell fate. Nature Materials 13, 645–652 (2014).

46. Balestrini, J.L., Chaudhry, S., Sarrazy, V., Koehler, A. & Hinz, B. The mechanical memory of lung myofibroblasts. Integrative Biology 4, 410–421 (2012).

47. Dunham, C., Havlioglu, N., Chamberlain, A., Lake, S. & Meyer, G. Adipose stem cells exhibit mechanical memory and reduce fibrotic contracture in a rat elbow injury model. The FASEB Journal 34, 12976–12990 (2020).

48. Scott, A.K. et al. Mechanical memory stored through epigenetic remodeling reduces cell therapeutic potential. Biophys J 122, 1428–1444 (2023).

49. Price, C.C., Mathur, J., Boerckel, J.D., Pathak, A. & Shenoy, V.B. Dynamic self-reinforcement of gene expression determines acquisition of cellular mechanical memory. Biophysical Journal 120, 5074–5089 (2021).

50. Moon, S.Y. et al. Cell contractility drives mechanical memory of oral squamous cell carcinoma. Mol Biol Cell, mbcE22070266 (2023).

51. Yang, J. et al. Targeting LOXL2 for cardiac interstitial fibrosis and heart failure treatment. Nat Commun 7, 13710 (2016).

52. Andres-Delgado, L. & Mercader, N. Interplay between cardiac function and heart development. Biochim Biophys Acta 1863, 1707–1716 (2016).

53. Johnson, P., Maxwell, D.J., Tynan, M.J. & Allan, L.D. Intracardiac pressures in the human fetus. Heart 84, 59–63 (2000).

54. McNally, E.M., Golbus, J.R. & Puckelwartz, M.J. Genetic mutations and mechanisms in dilated cardiomyopathy. J Clin Invest 123, 19–26 (2013).

55. Sanderson, J.E. Heart failure with a normal ejection fraction. Heart 93, 155–158 (2007).

56. Yamamoto, K. et al. Local neurohumoral regulation in the transition to isolated diastolic heart failure in hypertensive heart disease: absence of AT1 receptor downregulation and ‘overdrive’ of the endothelin system. Cardiovasc Res 46, 421–432 (2000).

57. Masuyama, T. et al. Evolving changes in Doppler mitral flow velocity pattern in rats with hypertensive hypertrophy. J Am Coll Cardiol 36, 2333–2338 (2000).

58. Roe, A.T. et al. Increased passive stiffness promotes diastolic dysfunction despite improved Ca2+ handling during left ventricular concentric hypertrophy. Cardiovasc Res 113, 1161–1172 (2017).

59. Ohmura, H. et al. Cardiomyocyte-specific transgenic expression of lysyl oxidase-like protein-1 induces cardiac hypertrophy in mice. Hypertens Res 35, 1063–1068 (2012).

60. Maitra, N., Flink, I.L., Bahl, J.J. & Morkin, E. Expression of alpha and beta integrins during terminal differentiation of cardiomyocytes. Cardiovasc Res 47, 715–725 (2000).

61. Terracio, L. et al. Expression of collagen binding integrins during cardiac development and hypertrophy. Circulation research 68, 734–744 (1991).

62. Brancaccio, M. et al. Differential onset of expression of α7 and β1D integrins during mouse heart and skeletal muscle development. Cell adhesion and communication 5, 193–205 (1998).

63. Nishiuchi, R. et al. Ligand-binding specificities of laminin-binding integrins: a comprehensive survey of laminin-integrin interactions using recombinant alpha3beta1, alpha6beta1, alpha7beta1 and alpha6beta4 integrins. Matrix Biol 25, 189–197 (2006).

64. Ziober, B.L. et al. Alternative extracellular and cytoplasmic domains of the integrin alpha 7 subunit are differentially expressed during development. J Biol Chem 268, 26773–26783 (1993).

65. Nawata, J. et al. Differential expression of alpha 1, alpha 3 and alpha 5 integrin subunits in acute and chronic stages of myocardial infarction in rats. Cardiovasc Res 43, 371–381 (1999).

66. Brancaccio, M. et al. Integrin signalling: the tug-of-war in heart hypertrophy. Cardiovasc Res 70, 422–433 (2006).

67. Liu, Y. et al. RNA-Seq identifies novel myocardial gene expression signatures of heart failure. Genomics 105, 83–89 (2015).

68. Manso, A.M. et al. Loss of mouse cardiomyocyte talin-1 and talin-2 leads to beta-1 integrin reduction, costameric instability, and dilated cardiomyopathy. Proc Natl Acad Sci U S A 114, E6250–E6259 (2017).

69. Manso, A.M. et al. Talin1 has unique expression versus talin 2 in the heart and modifies the hypertrophic response to pressure overload. J Biol Chem 288, 4252–4264 (2013).

70. Bogatan, S. et al. Talin Is Required Continuously for Cardiomyocyte Remodeling during Heart Growth in Drosophila. PLOS ONE 10, e0131238 (2015).

71. Goult, B.T. The Mechanical Basis of Memory - the MeshCODE Theory. Front Mol Neurosci 14, 592951 (2021).

72. Arias-Salgado, E.G. et al. Src kinase activation by direct interaction with the integrin beta cytoplasmic domain. Proc Natl Acad Sci U S A 100, 13298–13302 (2003).

73. Roca-Cusachs, P., Iskratsch, T. & Sheetz, M.P. Finding the weakest link: exploring integrin-mediated mechanical molecular pathways. J Cell Sci 125, 3025–3038 (2012).

74. Li, G. et al. Full activity of the deleted in liver cancer 1 (DLC1) tumor suppressor depends on an LD-like motif that binds talin and focal adhesion kinase (FAK). Proceedings of the National Academy of Sciences 108, 17129–17134 (2011).

75. Su, B. & Wu, J. Phosphorylation of RIAM Activates Its Adaptor Function in Mediating Integrin Signaling. J Cell Signal 2, 103–110 (2021).

76. Kanchanawong, P. et al. Nanoscale architecture of integrin-based cell adhesions. Nature 468, 580–584 (2010).

77. Sah, V.P. et al. Cardiac-specific overexpression of RhoA results in sinus and atrioventricular nodal dysfunction and contractile failure. J Clin Invest 103, 1627–1634 (1999).

78. Gunawan, M. et al. The methyltransferase Ezh2 controls cell adhesion and migration through direct methylation of the extranuclear regulatory protein talin. Nat Immunol 16, 505–516 (2015).

79. Gough, R.E. et al. Talin mechanosensitivity is modulated by a direct interaction with cyclin-dependent kinase-1. J Biol Chem 297, 100837 (2021).

80. Tabdanov, E. et al. Micropatterning of TCR and LFA-1 ligands reveals complementary effects on cytoskeleton mechanics in T cells. Integr Biol (Camb) 7, 1272–1284 (2015).

81. Hawkes, W. et al. Regulation of cardiomyocyte adhesion and mechanosignalling through distinct nanoscale behaviour of integrin ligands mimicking healthy or fibrotic extracellular matrix. Philos Trans R Soc Lond B Biol Sci 377, 20220021 (2022).

82. Levitt, J.A. et al. Quantitative real-time imaging of intracellular FRET biosensor dynamics using rapid multi-beam confocal FLIM. Sci Rep 10, 5146 (2020).

83. Tucker, N.R. et al. Transcriptional and Cellular Diversity of the Human Heart. Circulation 142, 466–482 (2020).

84. Jiang, H. et al. Functional analysis of a gene-edited mouse model to gain insights into the disease mechanisms of a titin missense variant. Basic Res Cardiol 116, 14 (2021).

