## Supplementary Figures and Table for "Cardiomyocyte mechanical memory is regulated through the talin interactome and DLC1 dependent regulation of RhoA"

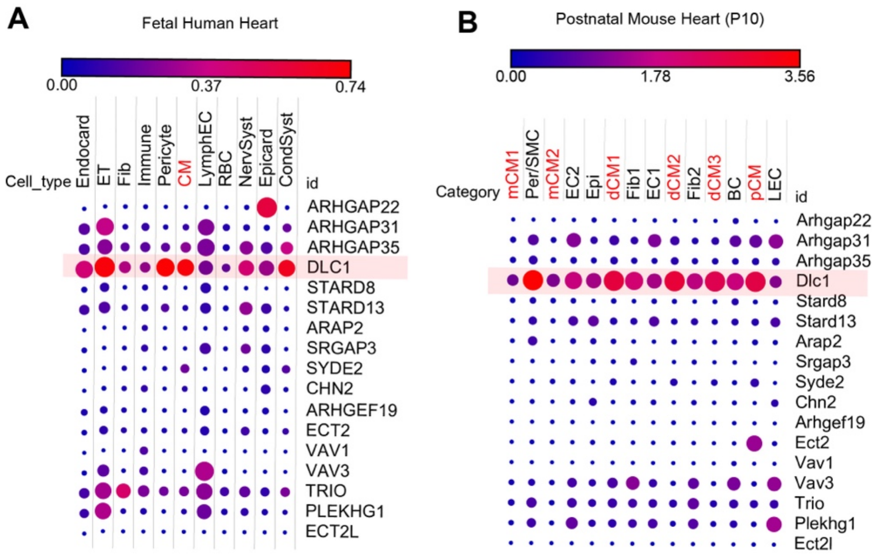


**Supplementary Figure S1: DLC1 is highly expressed in fetal human hearts and postnatal mouse hearts.** Expression of adhesion localised RhoA, Rac1 and CDC42 GAPs and GEFs (as identified by Müller et al [31]) was analysed using the broad institute single cell portal in **A)** single cell RNA-Seq data of a normal perinatal human heart (13569 cells, gestational age 83 days)[32] **B)** single nuclei RNA-Seq data from postnatal mouse hearts (7760 nuclei from 3 wild type mouse hearts at P10)[34] .

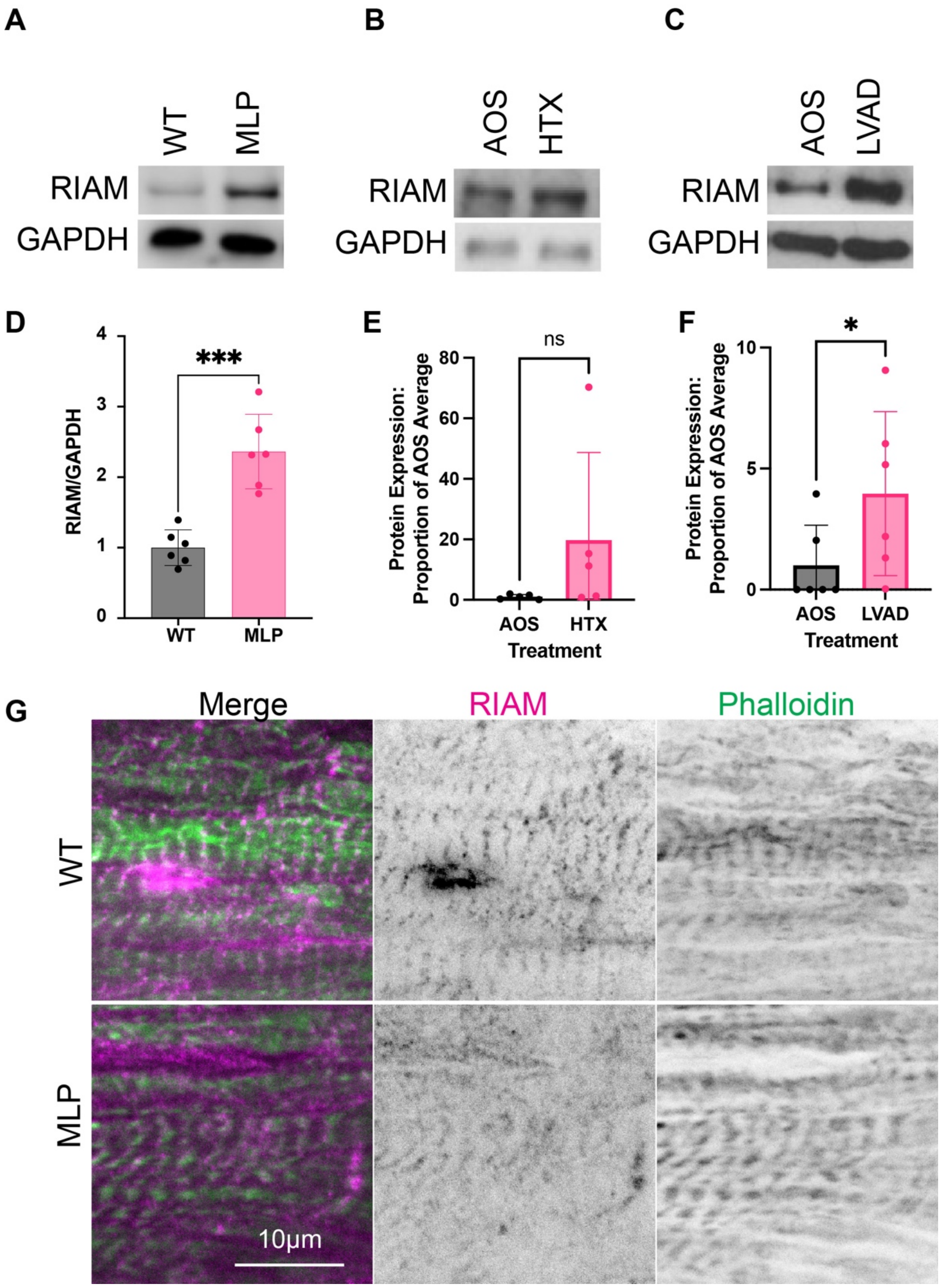


**Supplementary Figure S2: RIAM expression is increased in heart disease. A)** RIAM is upregulated in MLP knockout hearts; **B)** in failing explanted human hearts at time of explant (HTX) compared to aortic stenosis with preserved ejection fraction as control (AOS, since no healthy human heart samples were available); **C)** in failing hearts at time of implant of left ventricular assistance device (LVAD), compared to AOS; **D-F)** Quantification of (A), (B) and (C), respectively; **G)** Immunostaining of mouse heart sections (left ventricle) show costameric RIAM staining in both wild type and MLP knockout mouse hearts. *$p<0.0332$, ***$p<0.0002$; p-values from unpaired t-test.

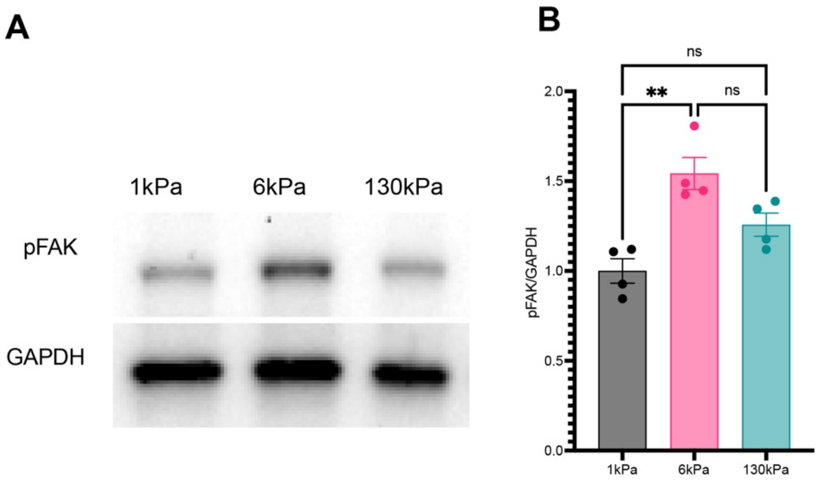


**Supplementary Figure S3: FAK activity is increased at 6kPa.** Neonatal rat cardiomyocytes were cultured on PDMS with the indicated stiffness, before lysis and blotting with pY397 FAK and GAPDH antibodies. *p<0.0332, **p<0.0021, ***p<0.0002, ****p<0.0001; p-values from one-way ANOVA with Dunnett correction for multiple comparisons.

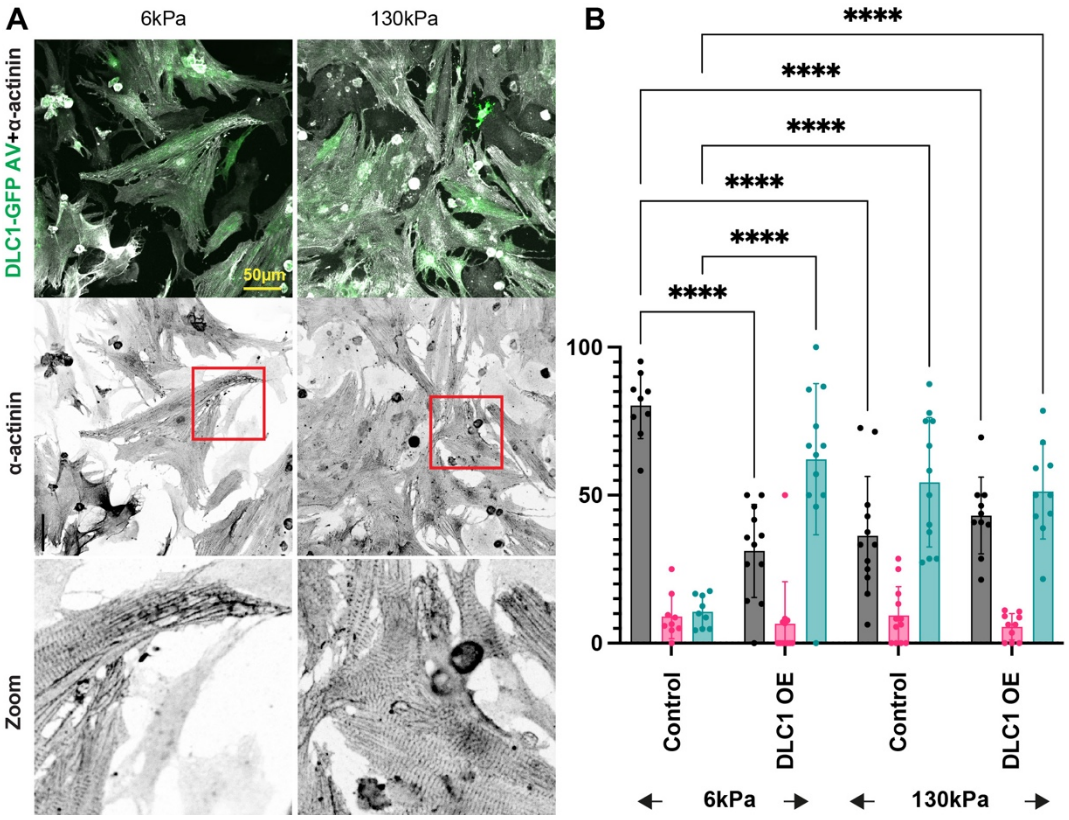


**Supplementary Figure S4: DLC1 overexpression leads to increased stress-fibre like structures.** Neonatal rat cardiomyocytes were cultured on PDMS with the indicated stiffness. Infection with a DLC1-GFP adenoviral vector (DLC1 OE) results in increased stress-fibre like structures. *p<0.0332, **p<0.0021, ***p<0.0002, ****p<0.0001; p-values from one-way ANOVA with Dunnett correction for multiple comparisons.

**Supplementary Tables:**

**Supplementary Table 1:**

Clinical data of patients providing cardiac tissue for Western blots. HTx – heart transplant, AoS – aortic stenosis, LVAD – left ventricular assist device, DM – diabetes mellitus, NIDDM – non-insulin-dependent diabetes, IDDM – insulin dependent diabetes mellitus, COPD – chronic obstructive pulmonary disease, CAD – coronary artery disease, MI – myocardial infarction, EF – ejection fraction, SR – sinus rhythm, AF – atrial fibrillation , ICD – implantable cardioverter defibrillator, ECMO – extracorporeall membrane oxygenation, PMK/ICD – pacemaker/ implantable cardioverter, PTCA – percutaneous transluminal coronary angioplasty, Cx – circumflex artery, CABG – coronary artery bypass graft, AVR - aortic Valve Replacement

| No. | Type | Sample region | Age at intervention | Hypertension | DM (0 = no, 1 = NIDDM, 2 = IDDM) | Dyslipidemia (0 = no, 1 = yes) | Smoke (0 = no, 1 = active smoker, 2 = former) | COPD | CAD | Previous MI | EF | Heart rhythm | Previous Cardiac Surgery | Valvular Disease | Cardiomyopathy | Operation |
|---|---|---|---|---|---|---|---|---|---|---|---|---|---|---|---|---|
| 2 | HTx | LV | 38 | 0 | 0 | 0 | 0 | 0 | 0 | 0 | 27 | SR | ICD implantation, ECMO | Severe Mitral Regurgitation | DCM | Heart Transplant |
| 3 | HTx | LV | 63 | 0 | 0 | 0 | 0 | 0 | 0 | 0 | 30 | AF | ICD implantation, Mitral Valve Replacement | 0 | DCM | Heart Transplant |
| 4 | HTx | LV | 52 | 0 | 0 | 0 | 0 | 0 | 0 | 0 | 15 | SR | ICD implantation, ECMO | 0 | ACM | Heart Transplant |
| 5 | HTx | LV | 66 | 1 | 0 | 0 | 0 | 0 | 0 | 0 | 15 | SR | ICD implantation | Moderate Mitral Regurgitation - Moderate Tricuspid Regurgitation | DCM | Heart Transplant |
| 6 | HTx | LV | 64 | 1 | 0 | 0 | 2 | 0 | 0 | 0 | 20 | SR | ICD implantation | Severe Mitral Regurgitation | DCM | Heart Transplant |
| 8 | HTx | LV | 68 | 0 | 0 | 0 | 2 | 0 | 0 | 0 | 15 | AF | ICD implantation | 0 | DCM | Heart Transplant |
| 9 | LVAD | LV apex | 54 | 1 | 0 | 1 | 2 | 1 | 1 | 1 | 20 | SR | ICD implantation | 0 | IHD | LVAD |
| 10 | LVAD | LV apex | 57 | 1 | 0 | 0 | 0 | 0 | 1 | 1 | 18 | SR | ICD implantation | 0 | IHD | LVAD |
| 11 | LVAD | LV apex | 64 | 1 | 0 | 0 | 0 | 0 | 1 | 1 | 19 | SR | ICD implantation | 0 | IHD | LVAD |
| 12 | LVAD | LV apex | 59 | 0 | 0 | 0 | 0 | 0 | 0 | 0 | 19 | SR | ICD implantation | 0 | DCM | LVAD |
| 13 | LVAD | LV apex | 48 | 1 | 1 | 1 | 0 | 0 | 1 | 1 | 20 | SR | ICD implantation | 0 | IHD | LVAD |
| 17 | LVAD | LV apex | 66 | 1 | 1 | 1 | 0 | 0 | 0 | 0 | 36 | Paroxismal AF | ICD implantation | 0 | DCM | LVAD |
| 18 | AoS | LV | 71 | 1 | 0 | 1 | 2 | 0 | 1 | 1 | 27 | SR | PMK/ICD implantation | Severe Aortic Stenosis | 0 | AVR + 1 CABG |
| 23 | AoS | LV | 64 | 1 | 0 | 1 | 0 | 0 | 1 | 0 | 55 | SR | PTCA on Cx | Severe Aortic Stenosis | 0 | AVR |
| 24 | AoS | LV | 56 | 1 | 0 | 1 | 0 | 0 | 0 | 0 | 71 | SR | 0 | Severe Aortic Stenosis | 0 | AVR + subaortic membrane removal |
| 28 | AoS | LV | 72 | 1 | 0 | 1 | 0 | 0 | 1 | 0 | 57 | SR | 0 | Severe Aortic Stenosis | 0 | AVR + 1 CABG |
| 32 | AoS | LV | 78 | 1 | 0 | 1 | 0 | 0 | 1 | 0 | 55 | SR | 0 | Severe Aortic Stenosis | 0 | AVR + 2 CABG |
| 33 | AoS | LV | 82 | 1 | 0 | 0 | 0 | 0 | 1 | 0 | 65 | SR | 0 | Severe Aortic Stenosis | 0 | AVR + 2 CABG |
